# RippleNet: A Recurrent Neural Network for Sharp Wave Ripple (SPW-R) Detection

**DOI:** 10.1101/2020.05.11.087874

**Authors:** Espen Hagen, Anna R. Chambers, Gaute T. Einevoll, Klas H. Pettersen, Rune Enger, Alexander J. Stasik

## Abstract

Hippocampal sharp wave ripples (SPW-R) have been identified as key bio-markers of important brain functions such as memory consolidation and decision making. SPW-R detection typically relies on hand-crafted feature extraction, and laborious manual curation is often required. In this multidisciplinary study, we propose a novel, self-improving artificial intelligence (AI) method in the form of deep Recurrent Neural Networks (RNN) with Long Short-Term memory (LSTM) layers that can learn features of SPW-R events from raw, labeled input data. The algorithm is trained using supervised learning on hand-curated data sets with SPW-R events. The input to the algorithm is the local field potential (LFP), the low-frequency part of extracellularly recorded electric potentials from the CA1 region of the hippocampus. The output prediction can be interpreted as the time-varying probability of SPW-R events for the duration of the input. A simple thresholding applied to the output probabilities is found to identify times of events with high precision. The reference implementation of the algorithm, named ‘RippleNet’, is open source, freely available, and implemented using a common open-source framework for neural networks (`tensorflow.keras`) and can be easily incorporated into existing data analysis workflows for processing experimental data.

## 1 Introduction

### Experimental background

Sharp wave ripples (SPW-R) are highly synchronous, fast oscillations observed in the CA1 region of the hippocampus of mammals, and are linked to mechanisms that play important roles in memory function (Buzsáki 2015). The oscillations associated with SPW-Rs are typically observed in the local field potential (LFP), which is the low-frequency (≲ 300 Hz) part of extracellularly recorded electric potentials measured in neural tissue. SPW-Rs arise in sleep and resting states and consist of large amplitude deflections of the local field potential (LFP) signal originating in the hippocampal CA3 region (‘sharp waves’), that can elicit fast oscillations in the hippocampal CA1 region (‘ripples’). Excitatory output from the CA1 region during ripples encodes sequences of neuronal activation of awake experiences, that reaches wide areas of the cortex as well as subcortical nuclei. For a comprehensive review on SPW-Rs, their origin and function, see for example Buzsáki 2015.

The SPW-R oscillations are observed above the cortical *γ*-band frequencies (30 − 90 Hz) (Silva 2013) of the LFP, and lie between 160 − 180 Hz in mice (Buzsáki, Logothetis, and Singer 2013; Buzsáki et al. 2003), and between 130 − 160 Hz in rats (Buzsáki, Logothetis, and Singer 2013; Buzsaki et al. 1992; John O’Keefe 1978). Features of one such example SPW-R event recorded using the experimental setup in Figure 1A,B are illustrated in Figure 1C. The wide-band LFP (top panel) contains a transient oscillation in its 150-250 Hz range (middle panel), also evident in the time-frequency resolved LFP spectrogram (bottom panel).

**Figure 1:**
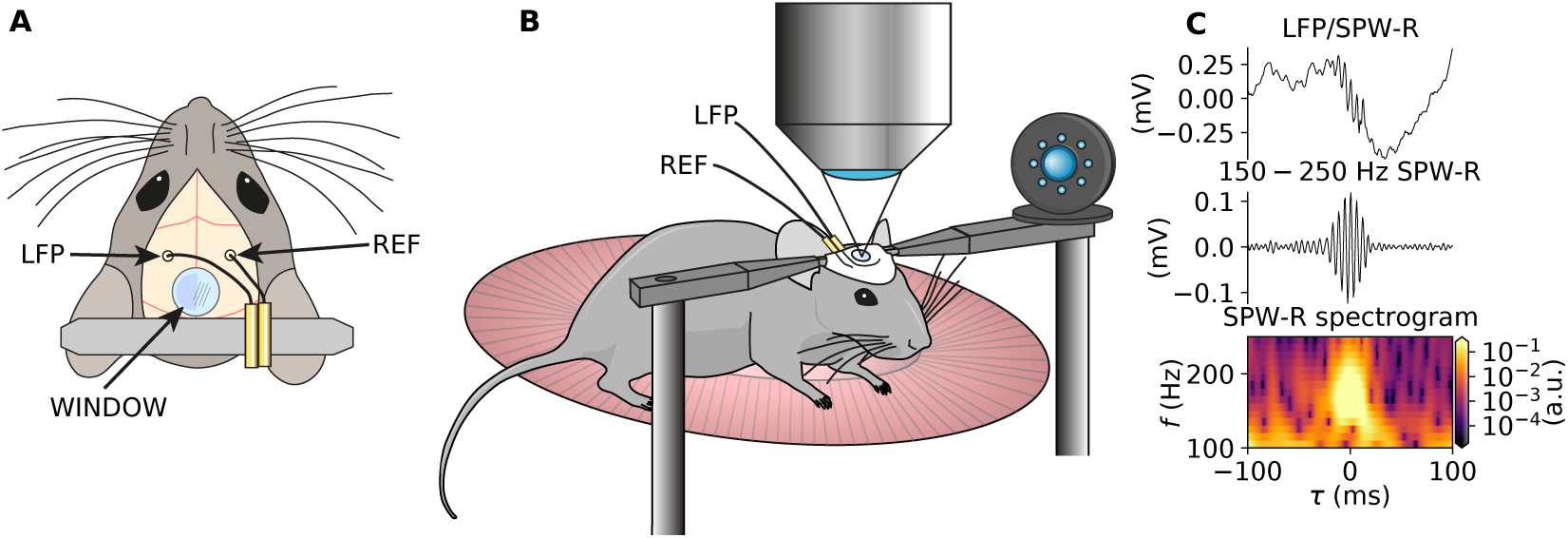
Experimental setup. A set of recordings used for this study were acquired concomitant to two-photon microscopy from head-fixed mice. A: Mice were prepared with a single electrode in the the hippocampal CA1 region and a contralateral reference electrode, chronic glass window for two-photon microscopy and a headbar for head-fixation. B: LFP recordings were mostly recorded concomitant to two-photon microscopy in head-fixed mice. C: Example of a single detected SPW-R. Top panel: raw LFP data; Middle panel: bandpass filtered LFP signal (150-250 Hz); Bottom panel: LFP spectrogram.

### SPW-R detection

The fractions of neurons in hippocampal regions CA1 and CA3 which are active during different SPW-R events vary greatly, and the number of small and medium-sized events outnumber large, highly synchronous events (Csicsvari et al. 1999a; Buzsáki 2015). Hence, the resulting distributions of SPW-R power are skewed as the synchrony between neurons throughout the network (i.e., correlations) greatly affects the LFP power in the SPW-R band (Csicsvari et al. 2000; Schomburg et al. 2012; Buzsáki 2015; Hagen et al. 2016). Consequently, defining a fixed threshold for SPW-R detection based on e.g., the power or envelope of the LFP in a chosen frequency band (Csicsvari et al. 1999a; Csicsvari et al. 1999b; Einevoll et al. 2013; Ramirez-Villegas, Logothetis, and Besserve 2015; Norman et al. 2019; Tingley and Buzsáki 2020) remains heuristic.

Different existing and novel real time algorithms for SPW-R detection were reviewed and tested on synthesized datas by Sethi and Kemere 2014. Such methods are applied with band-pass filtered LFP data, commonly in the 150 − 250 Hz range, and may incorporate adaptive thresholding (Fritsch, Ibanez, and Parrilla 1999; Jadhav et al. 2012).

### Deep learning

Recent years have seen a surge in different supervised and unsupervised learning algorithms, propelled by hardware acceleration, better training datasets and the advent of deep convolutional neural networks (CNN) in image classification and segmentation tasks (see e.g., Le-Cun, Bengio, and Hinton 2015; Rawat and Wang 2017). Deep CNNs are, however, not yet as commonplace for time series classification tasks (Fawaz et al. 2019). Unlike traditional neural networks (NNs) and CNNs which typically employ a feed-forward hierarchical propagation of activation across layers, *recurrent neural networks* (RNN) have feedback connections, and is suitable for sequential data such as speech and written text. One architecture of RNNs is Long Short-Term Memory (LSTM) RNNs (Hochreiter and Schmidhuber 1997), capable of classifying, processing and predicting events in time-series data, even in the presence of lags of unknown duration.

In speech recognition, deep RNNs with multiple stacked LSTM layers have been successful in classifying phonemes (Graves, Mohamed, and Hinton 2013). Graves, Mohamed, and Hinton 2013 also found bidirectional LSTM RNNs to improve classification performance over unidirectional LSTM RNNs, which can only account for past context. The present context of SPW-R detection is analogous and amounts to recognition of a single phoneme or word in a temporal sequential signal such as sound.

### Summary of present work

Here, inspired by RNNs shown to be successful on speech-recognition tasks, we propose the utilization of LSTM-based RNNs for the automated detection of SPW-R events in continuous LFP data. Our open-source implementation, RippleNet, is built with a combination of convolutional, (bidirectional) LSTM and dense output layers with non-linear activation functions. RippleNet accepts raw LFP traces of arbitrary length as input, and omits the typical SPW-R detection steps of band-pass filtering the input LFP as well as manual feature extractions such as computing the signal envelope via the Hilbert transform or time-frequency resolved spectrograms. Using training data with labeled SPW-R events in real-world datasets from different sources we trained RippleNet to predict a continuous signal representing the time varying probability of SPW-R event in the input. A simple search of local peaks above a fixed threshold can then be applied with the output probabilities, and is here shown to yield accurate predictions of the time of SPW-R events in separate validation and test datasets, with low prediction rates of false positive (FP) and false negative (FN) events. RippleNet also runs faster than real-time on typical CPUs, and even faster on graphical processing units (GPU).

RippleNet’s implementation differs from Zuo et al. 2019, one of very few deep CNN based algorithms specifically designed for detection of high-frequency oscillations (HFO), that is, epileptogenic zone seizures in intracranial electroencephalogram (iEEG) recordings, in that (1) explicit conversion of 1D input sequences with multiple rows into gray-scale images are avoided; (2) normalization of the input to zero mean and standard deviation to unity is not required; (3) input segments can be of arbitrary length (i.e., continuous) but segments within single batches have to be of the same length; (4) a fairly low number of parameters are trainable which may reduce overfitting; and (5) its outputs are continuous signals that represent the time varying probability of an SPW-R event at all time points of the inputs, in contrast to classifying whether or not a HFO class occurs in each fixed-size input segment.

### Manuscript structure

This paper is organized as follows: In Methods we detail the acquisition of experimental LFP data, labeling of SPW-R events, preprocessing steps, numerical analysis, and the technical implementation of RippleNet. In Results we evaluate the performance of RippleNet during training and application on separate validation and test sets. In Discussion we discuss the possible consequences, extensions and other applications of this work.

## 2 Methods

### 2.1 Experimental data

#### 2.1.1 Mouse data

Male and female mice (C57Bl/6J; Janvier labs) underwent LFP electrode implant surgery at approximately 10-14 weeks of age. All mice had previously been implanted 2-3 weeks earlier with a custom made titanium headbar glued to the skull and covered with a dental cement cap. For electrode implant surgery, mice were anesthetized with isoflurane (3 induction, 1.5 maintenance) with body temperature maintained at 37 degrees Celsius. Burr holes were drilled for the LFP electrode and reference electrode over the dorsal CA1 region of the hippocampus (A/P −2 mm, M/L 2 mm) and contralateral primary somatosensory cortex (A/P – 0.5 mm, M/L 3 mm), respectively. Silver wire electrodes (1.25 mm diameter, insulated, GoodFellow) were lowered to a depth of 0.8 mm for dorsal CA1. The reference electrode was implanted at the brain surface. Mice were allowed to recover from isoflurane anesthesia while headfixed for at least 15 minutes, and electrode placement was confirmed by monitoring the LFP signal online. Electrodes were affixed to the headbar with cyanoacrylate glue and a thin layer of dental cement.

All procedures were approved by the Norwegian Food Safety Authority (project: FOTS 19129). The experiments were performed in accordance with the Norwegian Animal Welfare Act and the European Convention for the Protection of Vertebrate Animals used for Experimental and Other Scientific Purposes.

LFP recordings were band-pass filtered (0.1-1000 Hz) and amplified (1000x) with a DAM50 differential amplifier (World Precision Instruments Inc). Line noise was removed using a HumBug 50/60 Hz Noise Eliminator (Quest Scientific Inc). For experiments, mice were headfixed under a two-photon microscope objective after brief isoflurane anesthesia. They were given at least 15 minutes to recover from anesthesia before recordings were taken. In most cases, LFP recordings were performed concurrently with two-photon calcium imaging through a chronic cranial window over the retrosplenial cortex, in 10 minute sessions. During recordings, mice were able to walk freely on a disc equipped with a rotary encoder to record locomotion, grooming and postural adjustments. Experiments were performed in the dark. LFP and rotary encoder signals were acquired at 20 kHz and downsampled to a final sampling frequency *f*_s_ = 2500 Hz in LabView (National Instruments). The LFP signals were saved in units of millivolts (mV).

The different animals, number of sessions, total recording durations and number of SPW-R events are listed in Table 1.

**Table 1:** Summary of data acquisition, and extracted training, validation and test data.

| Animals and sessions |  |  |  |
| --- | --- | --- | --- |
| mouse ID | # sessions | duration (s) | # SPW-R |
| 4028 | 4 | 2412 | 425 |
| 4029 | 1 | 603 | 86 |
| 4030 | 4 | 2412 | 658 |
| 4031 | 2 | 1206 | 176 |
| 4046 | 4 | 2410 | 596 |
| 4104 | 1 | 603 | 181 |
| 4105 | 3 | 1809 | 231 |
| 4106 | 4 | 2412 | 382 |
| 4214 | 3 | 1812 | 491 |
| 4215 | 3 | 1812 | 440 |
| 6102 | 3 | 1812 | 467 |
| 6112 | 3 | 1812 | 376 |
| rat ID | # sessions | duration (s) | # SPW-R |
| 2 | 1 | 29283 | 4498 |
| 7 | 1 | 25575 | 3714 |
| 9 | 1 | 26434 | 3677 |
| Extracted datasets |  |  |  |
| $\mathbf{X}, \mathbf{y}$ | $n_{\text{SPW-R}}: \{4461, 4400\}$ (mouse and rat)<br>shape: $(n_{\text{SPW-R}}, 1250, 1)$ | | |
| $\mathbf{X}_{\text{train}}, \mathbf{y}_{\text{train}}$ | $n_{\text{train}}: \{4175, 4000\}$<br>shape: $(n_{\text{train}}, 1250, 1)$ | | |
| $\mathbf{X}_{\text{val}}, \mathbf{y}_{\text{val}}$ | $n_{\text{val}}: \{200, 200\}$<br>shape: $(n_{\text{val}}, 1250, 1)$ | | |
| $\mathbf{X}_{\text{test}}, \mathbf{y}_{\text{test}}$ | $n_{\text{test}}: \{1, 0\}$<br>shape: $(n_{\text{test}}, 753914, 1)$ | | |

### SPW-R detection procedure

Pre-processing for manual SPW-R detection was performed using MATLAB 2018^1^. The LFP signal was first band-pass filtered between 150 and 250 Hz using a digital filter `filtfilt`. The coefficients for the order 600 finite impulse response (FIR) filter were generated using the `fir1` function. The band-pass filtered LFP was then used to compute the absolute of the Hilbert transform of the data. The output was smoothed by convolving with a 1052-point Gaussian filter with *σ* = 40 ms using `gaussfilt` (James Conder 2020). The `findpeaks` function was used to find peaks which were 3 standard deviations above mean in a moving time window with duration 1 s. A minimum peak width at half height of 15 ms and a minimal peak distance of 25 ms were required, calculated based on data reported in Axmacher, Elger, and Fell 2008; Davidson, Kloosterman, and Wilson 2009; Caputi et al. 2012. Potential ripple locations where then manually inspected using the symmetric one second time window around it, based on the Hilbert transformation and the raw LFP signal.

#### 2.1.2 Rat data

To supplement the training and validation datasets containing SPW-R events that could be extracted from the in-house datas described above, we utilized publicly available datasets from the Buzsaki lab webshare^2^ (Petersen, Hernandez, and Buzsáki 2018). The datas were obtained in the adult rat (Long Evans) in awake and sleep states using chronically implanted probes with a total of 96 or 128 channels (Tingley and Buzsáki 2018; Tingley and Buzsáki 2020). The datasets were

- DT2/DT2_rPPC_rCCG_3612um_1360um_20160303_160303_084915,
- DT7/20170324_576um_144um_170324_123932,
- DT9/20170509_468um_36um_170509_103451.

The LFP signal of contacts located in CA1 was extracted and converted to units of mV, along with the corresponding times and durations of labeled CA1 SPW-R events. SPW-R events were identified and labeled as described in Tingley and Buzsáki 2020. We here defined SPW-R event times as the mean of onset and offset times. All events in awake and sleep states were extracted. The sampling frequency of the LFP data was here *f*_s_ = 1250 Hz.

The different animals, number of sessions, total recording durations and number of SPW-R events are summarized in Table 1.

### 2.2 Data preprocessing

The mouse LFP data were downsampled to a common sampling frequency *f*_s_ = 1250 Hz and temporal resolution Δ*t* = 1*/f*_s_. For temporal downsampling we used the `scipy.signal.decimate` function with default parameters. The rat LFP datasets were used as is. For visualization, we extracted the band-pass filtered LFP *ϕ*_BP_(*t*) from the LFP *ϕ*(*t*) using 2nd-order Butterworth filter coefficients computed with critical frequencies *f*_c_ ∈ {150, 250} Hz. Filters were applied to *ϕ*(*t*) using a zero phase shift, forward-backward filter implementation. Filter coefficients were computed using `scipy.signal.butter` and applied with `scipy.signal.filtfilt`.

#### 2.2.1 Wavelet spectrograms

To compute spectrograms of LFP data *ϕ*(*t*) we relied on the complex Morlet transform with parameters *ω* = 6, scaling factor *s* = 1 and lengths *M*_*f*_ = 2*sf*_s_*ω/f* for fundamental frequencies *f* ∈ {100, 110 …, 240, 250} Hz. The numbers *M*_*f*_ were rounded down to the nearest integer. The set of discrete wavelet coefficients for each frequency *f* were computed using the function `scipy.signal.morlet` as

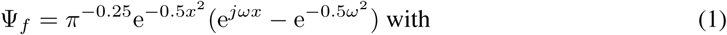

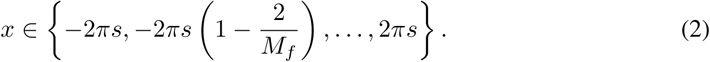

Each row of the spectrograms *S*(*t, f*) = [*S*_*f*_ (*t*)] were then computed for all frequencies in *f* as

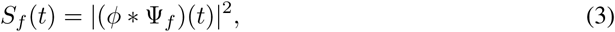

where the asterisk denotes a convolution. We used the discrete 1D implementation by `scipy.signal.convolution` in ‘full’ mode. To visualize the spectrograms, we employed a log-linear `matplotlib.cm.inferno` color map, with lower and upper limits determined as exp(*c*), where *c* is the 1% and 99% percentiles of log(*S*), respectively.

#### 2.2.2 Training, validation and test data

##### Input data

We chose to use the raw single-channel LFP data segments as input to the neural network algorithm, that is, by defining *X*(*t*) = [*ϕ*(*t*)]. For reasons related to the RNN implementation we defined each segment *X*(*t*) as shape (*n*_timesteps_, 1) arrays, even if we here work with single-channel LFP data.

##### One-hot encoding of SPW-R events

The train of *n* labeled times *t*^⟨*i*⟩^ of the SPW-R events in each continuous LFP data segment can be expressed mathematically as

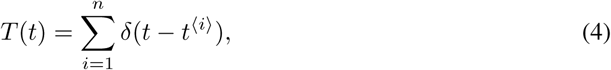

where *δ*(·) denotes the dirac delta distribution, and *i* the index of the event in a session. We then assumed that each SPW-R has a typical duration *D* = 50 ms on the interval [*t*^⟨*i*⟩^ − *D/*2, *t*^⟨*i*⟩^ +*D/*2). A binary ‘one-hot’ encoding vector for the SPW-R events *y*(*t*) was then computed as

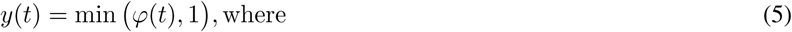

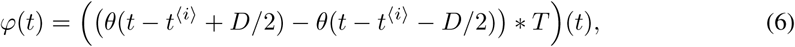

for the entire duration of each LFP segment. Here *θ*(·) denotes the Heaviside step function. The vector *y*(*t*) can be interpreted as the time-varying, binary probability *p* ∈ [0, 1] of an SPW-R occurring at any given time *t*.

##### Datasets

As the SPW-R occurrence in the data was *sparse* (that is, *y*(*t*) = 0 for most *t*), training the neural network on entire data segments of different durations is impractical. A likely training outcome is predicting *ŷ*(*t*) = 0 for all times *t* of the input. For each labeled SPW-R event we therefore extracted temporal segments of duration *T*_sample_ = 1000 ms from *X*(*t*) and *y*(*t*), that is, on the interval 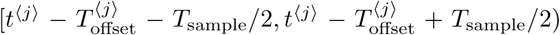. The offsets 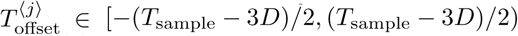 were randomly drawn for each event. The superscript ⟨*j*⟩ here denotes a sample indexed by *j* from any LFP recording session.

For the total number *n*_SPW − R_ of SPW-R samples across all animals and sessions, the shapes of the combined input and output dataset matrices **X** and **y** for training, validation and testing were both (*n*_SPW−R_, *T*_sample_*/*Δ*t*, 1).

All data entries except for a hold-out set were randomly reordered along their first axis, and then split into 2 separate file sets for validation and training, each of sizes summarized in table 1. The validation set was used to monitor loss during training and quantification of performance as detailed below. The hold-out test set constructed from the entire session of one animal (mouse 4029) was only utilized for final evaluation of the RNN after training and validation. For visualization purposes we also stored the corresponding segments of band-pass filtered LFP 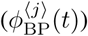 and spectrograms (*S*^⟨*j*⟩^(*t, f*)) for every labeled event.

In an effort to balance the set of features that can be learned by RippleNet from datas obtained mice and rat, we extracted a similar count of SPW-R events for training and validation from the rat data as for the mouse data.

### 2.3 RippleNet implementation

#### 2.3.1 Network description

The causal and non-causal RippleNet implementations, summarized schematically and with parameters in Table 2, consist of a Gaussian noise layer applied to the input, then one 1D convolutional layer followed by a dropout layer, followed by another 1D convolutional layer followed by batch-normalization, rectified-linear (ReLu) activation and dropout layers. The output of the last convolutional layer are consecutively fed to the first (bidirectional) LSTM layer followed by batch normalization and dropout. The final (bidirectional) LSTM layer is followed by a dropout, batch-normalization and a final dropout layer. Bidirectional layers are optionally applied using a wrapper function. The output is governed by a time-distributed layer wrapping a dense activation layer, which facilitates application of the dense layer to every temporal slice of the input. Hence, the output is a matrix of the same dimensionality as the input matrix.

**Table 2:** RippleNet neural network structure and parameters for both unidirectional and bidirectional variants.

| RippleNet architecture and settings |  |
| --- | --- |
| tf.keras.layer: | Parameters: |
| Input | shape: (None, 1) |
| ↓ |  |
| Gaussian | standard deviation: 0.001 |
| ↓ |  |
| Conv1D | filters: 20<br>kernel size: 11<br>strides: 1<br>use bias: False<br>padding: same<br>activation: None<br>parameters: 240 |
| Dropout | rate: 0.8 |
| ↓ |  |
| Conv1D | filters: 10<br>kernel size: 11<br>strides: 1<br>use bias: True<br>padding: same<br>activation: None<br>parameters: 2210 |
| BatchNormalization | parameters: 40 |
| Activation | activation: ReLu |
| Dropout | rate: 0.8 |
| ↓ |  |
| LSTM/<br>Bidirectional(LSTM) | units: 20/6<br>activation: tanh<br>recurrent activation: sigmoid<br>return sequences: True<br>parameters: 2480/816 |
| BatchNormalization | parameters: 80/48 |
| Dropout | rate: 0.8 |
| ↓ |  |
| LSTM/<br>Bidirectional(LSTM) | units: 20/6<br>activation: tanh<br>recurrent activation: sigmoid<br>return sequences: True<br>parameters: 3280/912 |
| Dropout | rate: 0.8 |
| BatchNormalization | parameters: 80/48 |
| Dropout | rate: 0.8 |
| ↓ |  |
| TimeDistributed(Dense) | nodes: 1<br>activation: sigmoid<br>parameters: 21/13 |

The Gaussian noise layer and dropout layers are only active during training in order to prevent overfitting of the training set, and inactive during validation and testing. The kernel weights of the convolutional, dense and LSTM layers are initialized with the Glorot uniform initializer. Recurrent connections in LSTM layers are initialized using the Orthogonal initializer. For optimization we chose the Adam algorithm which implements an adaptive stochastic gradient descent method (Kingma and Ba 2014). The settings for model compilation, optimization algorithm and model fitting are summarized in Table 3. For 3-fold cross-validation different RNN instances are initialized using different random seeds affecting initializers, dropout layers and optimization. This ensure replicable results on similar GPU hardware.

**Table 3:**
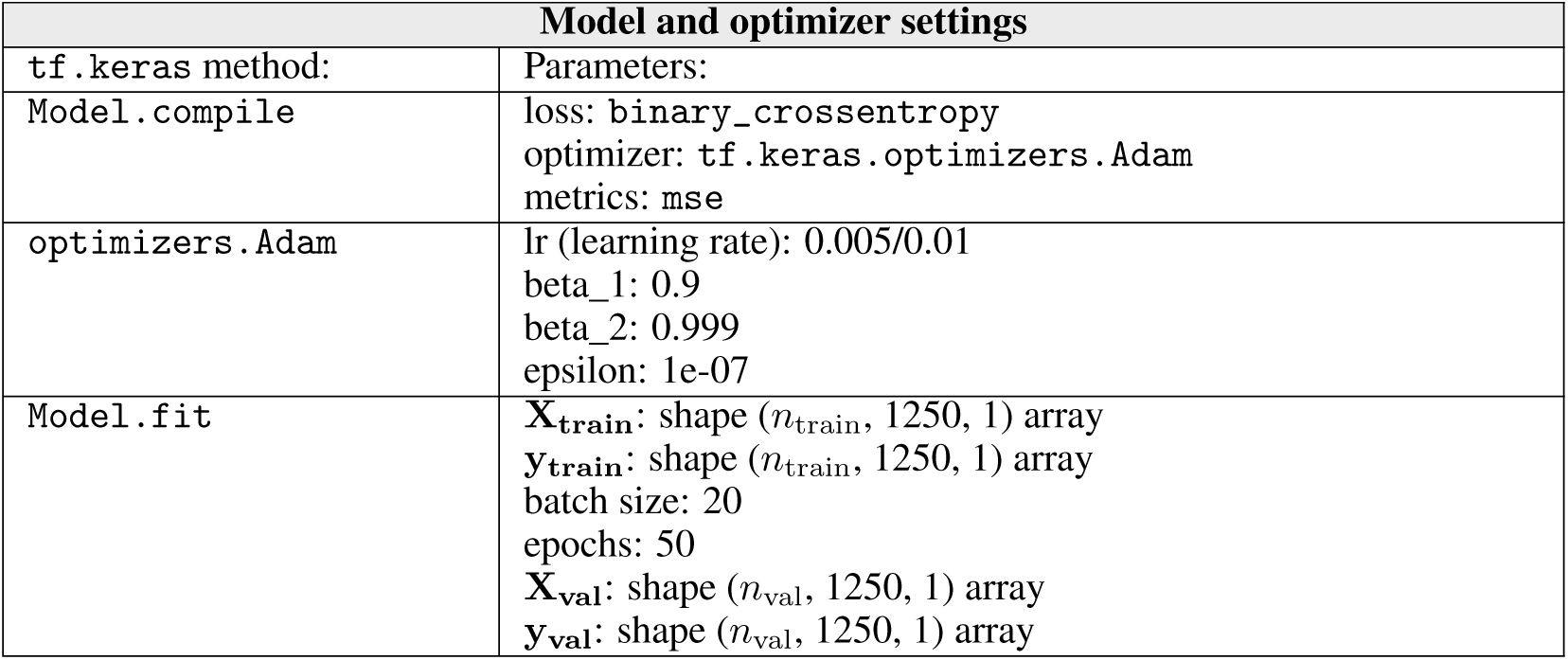
Summary of settings for model compilation, optimization and fitting of training data set.

Layer dimensions were hand tuned, with the aim of reducing the amount of trainable parameters and reducing evaluation times and overall training times, while maintaining achievable loss *J* and *MSE* reasonably low.

#### 2.3.2 Loss function and evaluation metric

For training the RNN we used the binary cross entropy loss function (`tf.keras.losses.BinaryCrossentropy`)

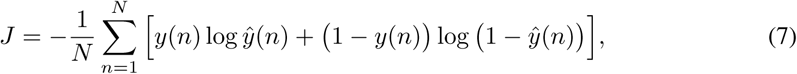

where *N* = *T*_sample_*/*Δ*t* is the number of temporal samples in the label array *y* and RNN prediction *ŷ*. To monitor training and validation performance of the RNN we used the mean squared error

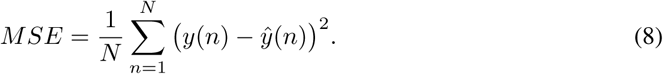

### 2.4 Data analysis

#### 2.4.1 Thresholding

The output *ŷ*^⟨*j*⟩^(*t*) ∈ (0, 1) of RippleNet is a discrete signal of same temporal duration and resolution as an input segment *X*^⟨*j*⟩^(*t*). The signal *ŷ*^⟨*j*⟩^(*t*) can be interpreted as the time-varying probability of an SPW-R ripple event. To extract time points of candidate ripple events, we ran the peak-finding algorithm implemented by `scipy.signal.find_peaks` using an initial threshold of 0.5, minimum peak inter-distance of 50 ms (same as *D*) and peak width of 20 ms. These parameters were set heuristically. Other parameters were left at their default values.

Further analyses of SPW-R detection performance were conducted by varying the threshold ∈ {0.1, 0.35, … 0.85, 0.95} and peak width ∈ {0, 5, …, 50} ms in a grid search, and assessing the influence on the metrics described next.

#### 2.4.2 Quantification of true and false predictions

On the validation and test data sets, we counted a true positive (*TP*) for the predicted time 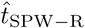 of an SPW-R event if 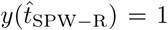, false positive (*FP*) if 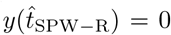 and false negative (*FN*) if no peaks above threshold in *ŷ*(*t*) were found in time intervals where *y*(*t*) = 1. *ŷ*(*t*) can be above threshold if FP predictions occur next to labeled SPW-R events and result in FN counts. Negative samples, where *y*(*t*) = 0 for all times spanned by the LFP samples, were not included in any of the training, validation or test sets. Hence, evaluation of true negative (*TN*) predictions were not performed. Note, however, that by construction, each sample *y*^⟨*j*⟩^(*t*) was equal to zero up to 95% of the time spanned by the sample, and that more than one SPW-R event may exist in each segment.

### 2.4.3 Precision, recall and *F*_1_ metrics

The following quantification metrics of SPW-R detection performance are used:

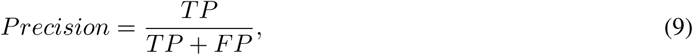

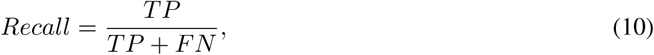

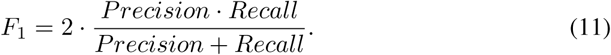

*Recall* is sometimes referred to as *True Positive Rate (TPR)* and *Sensitivity* in the literature (e.g., by Zuo et al. 2019). *Precision* is also known as *Positive Predictive Value (PPV).* The *F*_1_ score represents the harmonic mean of *Precision* and *Recall*. These metrics are all defined on the interval [0, 1], with 1 being best.

#### 2.4.4 Correlation analysis

To quantify the temporal agreement with labeled and predicted SPW-R event times, we compute the cross-correlation coefficients between predicted ripple event times 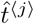 and labeled event times *t*^⟨*j*⟩^ as function of time lag *τ* as (Eggermont 2010, Eq. 5.10):

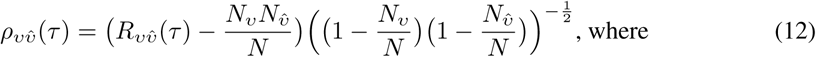

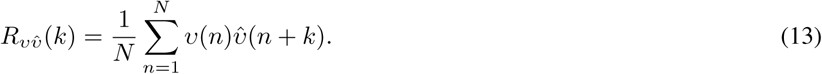

Here, *ν* and 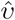 are the time binned *N*_*ν*_ and 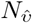 times of labeled and predicted SPW-R events using a bin width Δ = 2 ms, where 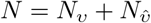.

#### 2.4.5 Signal energy

To quantify ‘strengths’ of ripples in the band-pass filtered LFP, we compute the signal energy (not to be confused with energy in physics) as

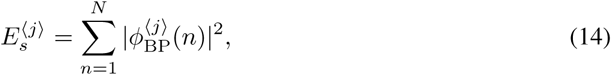

where *N* = 2*τ/*Δ*t* and *τ* ∈ [−100, 100] ms denotes time relative to the SPW-R event time.

### 2.5 Technical summary

The Python-based preprocessing and data extraction steps used Python^3^ (v3.6.10), jupyter-notebook^4^ (v6.0.3), numpy^5^ (v1.18.1, van der Walt, Colbert, and Varoquaux 2011), scipy^6^ (v1.4.1, Virtanen et al. 2020), h5py^7^ (v2.10.0, Collette et al. 2019), matplotlib^8^ (v3.2.1, Hunter 2007), pandas^9^ (v1.0.3, McKinney 2010) with the Anaconda Python Distribution^10^ (v4.8.3) running on a 13-inch 2016 Macbook Pro with macOS Mojave (v10.14.6).

The main training, analysis and visualization of performance of RippleNet was implemented and executed using Python (v3.6.9), jupyter-notebook (v5.2.2), numpy (v1.18.2), scipy (v1.4.1), h5py (v2.10.0), matplotlib (v3.2.1), seaborn^11^ (v0.10.1, Waskom et al. 2020), pandas (v1.0.3) and tensorflow^12^ (v2.1.0, Abadi et al. 2015) running on the Google Colaboratory portal^13^ using GPU hardware acceleration (using single Nvidia K80s, T4s, P4s or P100s cards).

## 3 Results

We here present our main findings and analysis of RippleNet, an automated, trainable recurrent neural network algorithm for detecting SPW-R events in single-channel LFP recordings.

### 3.1 Experimental data

Brain signals such as the LFP are characterized by low-frequency fluctuations, with spurious oscillatory events that may occur in different parts of the frequency spectrum. A few 1 s samples of hippocampus CA1 LFP, here used as validation data *X*^⟨*j*⟩^(*t*) ∈ **X**_val_ for the SPW-R detection algorithm, are shown in Figure 2A. Each sample contains at least one labeled SPW-R event at times marked by the green diamond symbols. The SPW-R events identified using a conventional method involving manual steps (cf. Methods), are hardly discernible by eye. They stand out, however, in the corresponding bandpass-filtered LFP signals 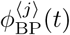 (panel B) and in the time-frequency resolved spectrograms *S*^⟨*j*⟩^(*t, f*) (panel C). Individual samples may also include potential SPW-R events that were not labeled. Events may have amplitudes of ∼0.1 mV in the filtered signal. Their durations are also short (≲ 100 ms).

**Figure 2:**
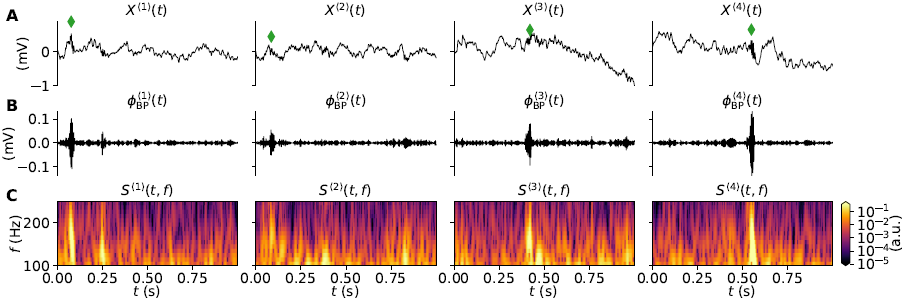
Snapshots of experimental data. A: Samples of raw LFP traces (*X*^⟨1⟩^(*t*), *X*^⟨2⟩^(*t*), …) with at least one SPW-R event. The green diamonds mark the times of the labeled events. Each column corresponds to samples *j* from the validation dataset. B: Bandpass-filtered LFP traces 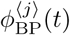. C: Wavelet spectrograms *S*^⟨*j*⟩^(*t, f*) computed from the LFP traces.

### 3.2 Training and validation of RippleNet

We next continue with 3-fold cross validation of different instances of RippleNet during and after training. We investigate both unidirectional (causal) and bidirectional (non-causal) variants of RippleNet, keeping the counts of trainable parameters about a factor two higher for the unidirectional variant (cf. Table 2). The computational load during training was approximately a factor two higher for the bidirectional variant. We observed ∼90 and ∼170 ms/step on Tesla P4 GPUs during training, respectively. The entire dataset (**X, y**) is split into separate training, validation and test sets with dimensions detailed in Table 1. Each RippleNet instance was initialized in each trial with different random seeds affecting initial weights, parameters etc., throughout the different model layers.

The training and validation loss *J* and *MSE* as function of training epoch are shown in Figure 3A,B, respectively. Continuous lines show training performance, while dashed lines show validation performance for different models. Models M1-3 are unidirectional, while models M4-6 are bidirectional. All models within each group, except M1, display similar and stable trajectories for their respective training sets. Validation loss and *MSE* are as expected inherently more variable across epochs, due to the smaller number of validation samples. The bidirectional variants perform consistently better than the unidirectional variants after just a few training epochs, both in terms of loss *J* and *MSE*. Validation loss *J* and *MSE* are reduced compared to training loss as noise and dropout layers are not active during validation. The different trajectories indicate no signs of over-fitting either to the training or validation sets.

**Figure 3:**
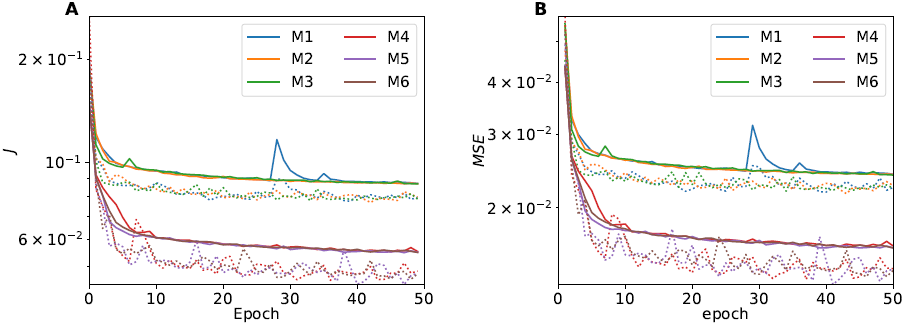
A: Training and validation loss *J* as function of training epoch for different RippleNet training runs, where initial network connection weights are instantiated using different random seeds. The continuous and dotted lines show loss on training and validation sets, respectively. B: *MSE* as function of training epoch.

#### 3.2.1 Validation set performance

Training and validation losses *J* and *MSE* only provide an indication of the ability to detect SPW-R events using the different models. First, in Figure 4 we visually compare a subset of predictions *ŷ*^⟨*j*⟩^(*t*) on LFP samples from a validation set (*X*^⟨*j*⟩^(*t*) ∈ **X**_val_(*t*)), to one-hot encoded SPW-R events *y*^⟨*j*⟩^(*t*) (see Methods). Here, all models produce predictions (panels C) with responses above the detection threshold for labeled events, but spurious threshold crossings may occur elsewhere.

**Figure 4:**
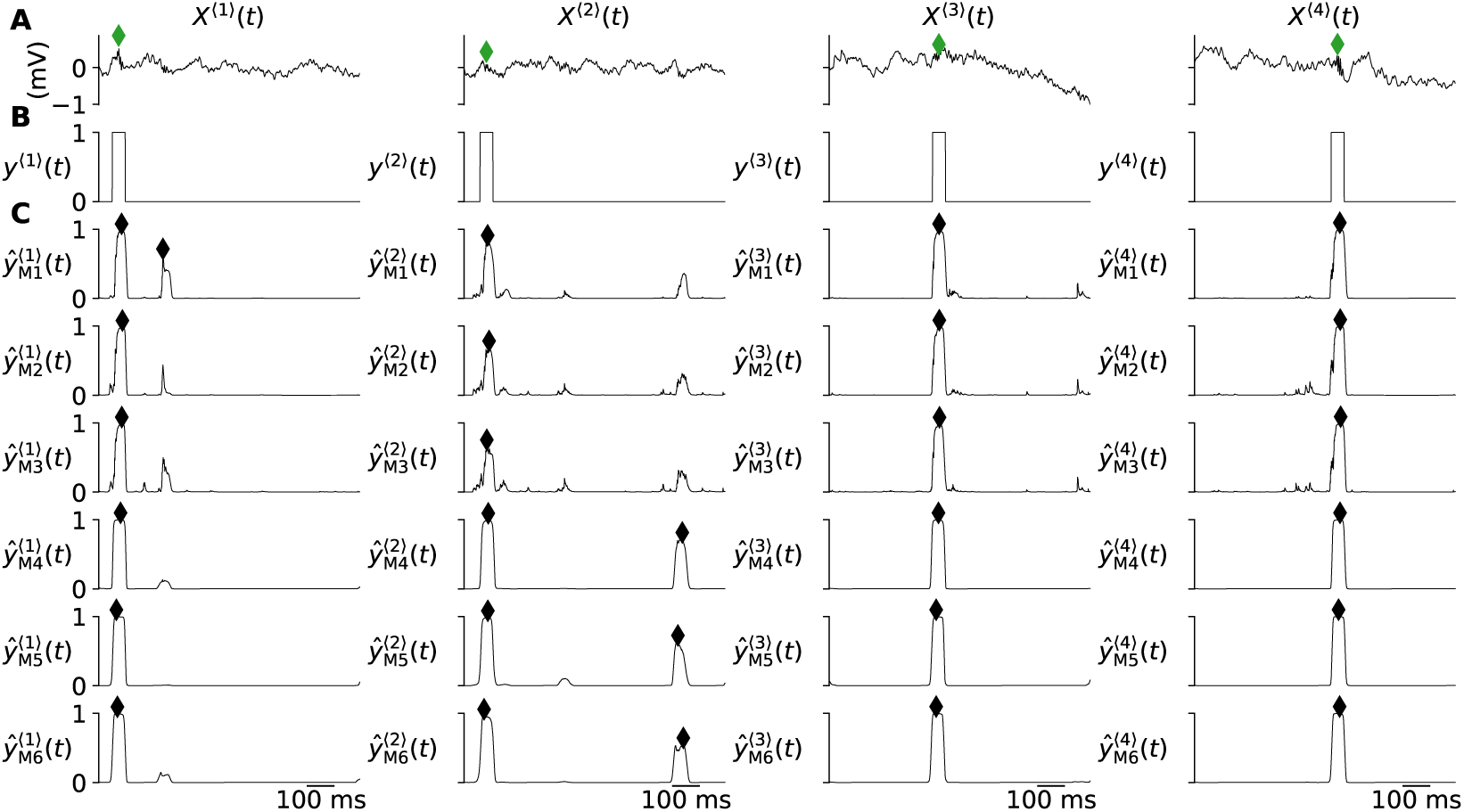
Comparison of RippleNet predictions on samples from the same test set. Each column corresponds to different LFP samples *X*^⟨*j*⟩^ shown at the top. A: Input LFP samples *X*^⟨*j*⟩^. The diamonds mark the labeled times of SPW-R events. B: Label vectors *y*^⟨*j*⟩^(*t*). C: Predictions *ŷ*^⟨*j*⟩^(*t*) made by the different RippleNet instances. SPW-R events found by the peak-finding algorithm are marked with diamond markers.

The non-causal bidirectional RippleNet variants (models M4-M6) produce output with notably less spurious fluctuations below threshold, when compared to the causal variants (M1-M3). These spurious fluctuations reflect the fact that signal power in the expected frequency range of SPW-R events do not vanish due to other ongoing neural processes, measurement noise etc. The bidirectional models do an overall better job at predicting the boxcar shapes of the one-hot encoded SPW-R events in panel B, owing to the fact that the full input time series are factored into their predictions.

We next quantify the different models’ performance in terms of counts of true positives (TP), false positives (FP) and false negative (FN) on the full validation set. Summarized in Table 4, trained models of each kind (uni-vs. bidirectional RippleNets) show similar numbers of TP events using the initial settings for the peak-finding algorithm when applied to the individual model predictions *ŷ*^⟨*j*⟩^(*t*). The counts of TPs are for all models generally in the same range, but total error counts (FP plus FN counts) are consistently higher for the unidirectional RippleNets compared to their bidirectional counterparts.

**Table 4:**
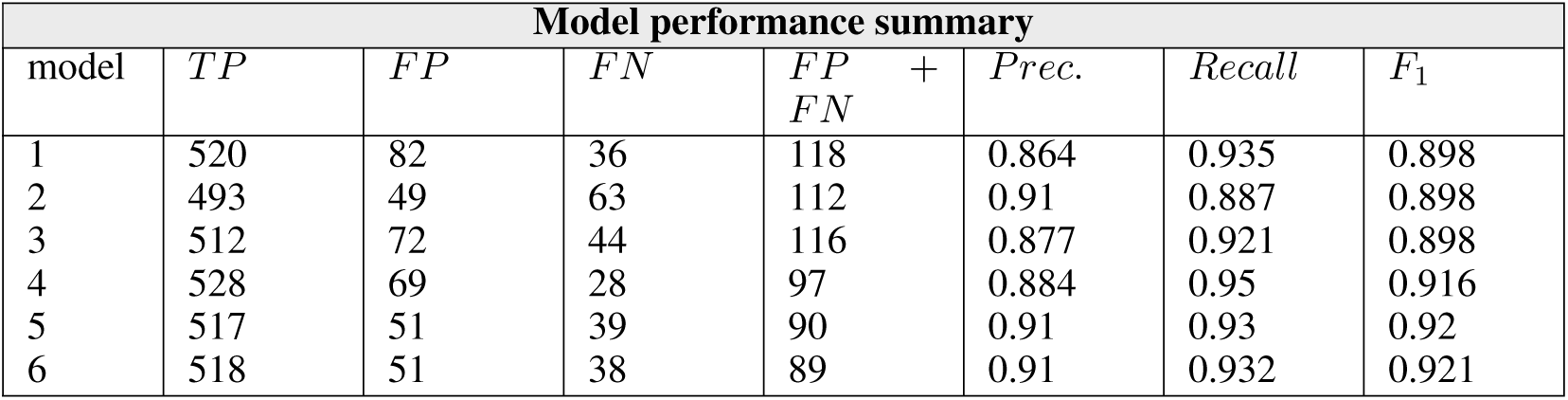
TP, FP, TN counts and performance metrics for RippleNet models on validation data.

In Table 4 we also compute the corresponding measures of performance from the TP, FP and FN counts. *Precision*, the ratio between TP predictions and total number of predictions, and *Recall*, the ratio between TP predictions and the sum of TP and FN predictions, are for most models around 0.9. Bidirectional models which show a better performance in terms of TP, FP and TN counts, and result in improved *F*_1_ scores of around 0.92.

#### 3.2.2 Effect of detection threshold and width parameters

The analysis above assumes fixed hyper-parameters for the peak-finding algorithm (cf. Methods) applied to the predictions by RippleNet instances on the validation set, including threshold, minimal peak interdistance and width (in units of time steps of size Δ*t*). We next hypothesize that the total error counts (FP+FN) can be minimized and correct prediction counts (TP) can be maximized using a hyper-parameter grid search, and therefore chose to optimize thresholds and widths for each network with respect to the *F*_1_-score. We keep the minimal peak interdistance the same as the boxcar filter width used to construct *y*(*t*). Summarized in Figure 5, the TP and FP counts for each model increased when lowering the threshold and width. FN counts increase for high threshold values and widths. Bidirectional models (M4-6) are less affected by the width setting compared to the unidirectional variants. The different models display different ‘sweet spots’ in terms of total number or errors (FP+FN). These counts are reflected in the calculated *Precision* and *Recall* values. The *F*_1_ space show for some models multiple local maxima. Here model 4 has the overall best performance, both in terms of least amounts of errors and highest *F*_1_ score. For further analysis and later application to a hidden test set we therefore choose that model, with detection threshold 0.7 and peak width of 0 time steps. In passing, we note that the other two bidirectional RippleNet instances achieve nearly similar levels of performance, while unidirectional variants have higher error counts.

**Figure 5:**
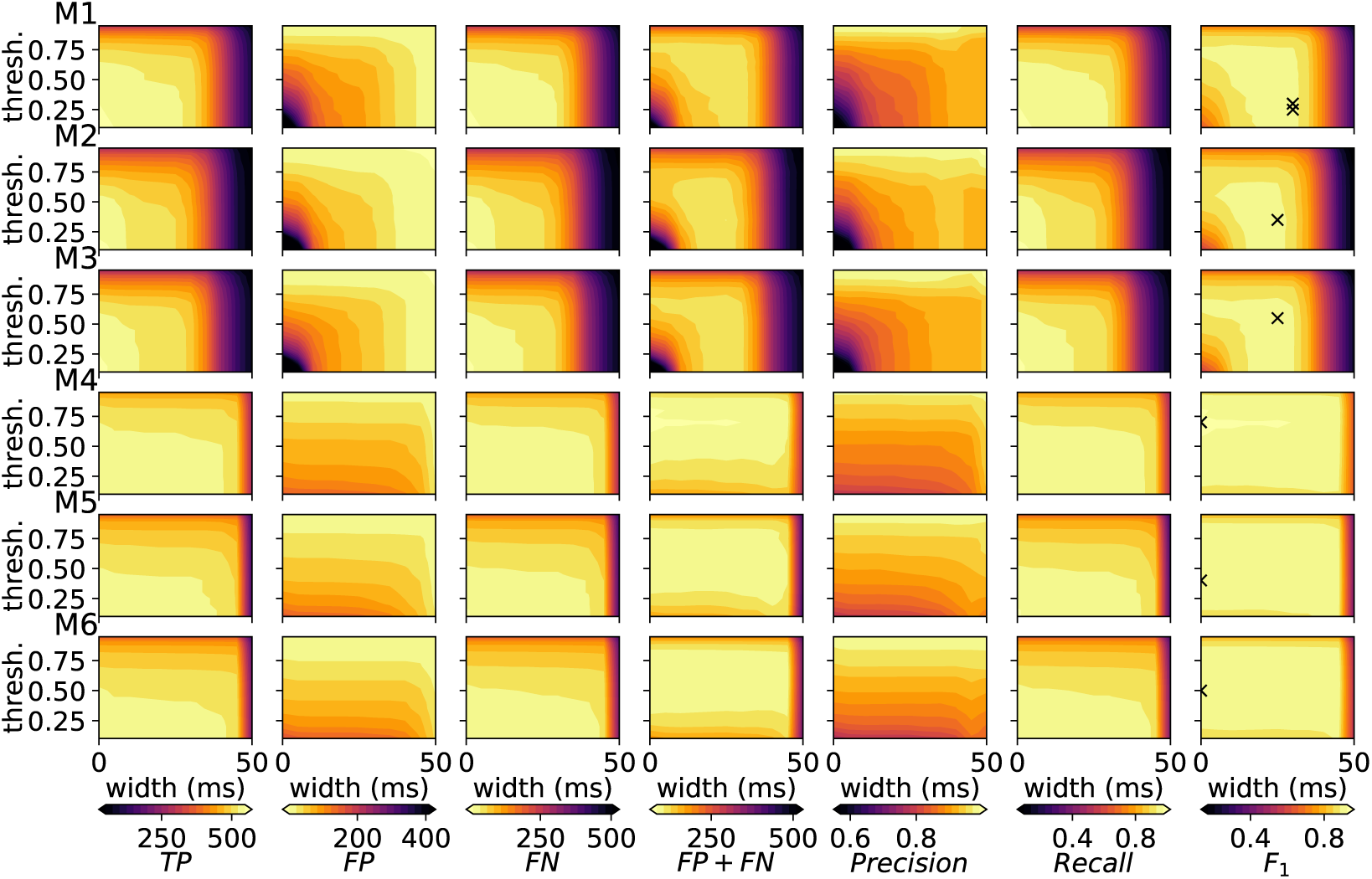
Effect of varying parameters threshold and width for the peak finding algorithm applied to predictions on the validation datas made by different RippleNet instances. Each row corresponds to different models and the columns to different metrics. Colorbars are shared among panels in each column. The cross hatches in the *F*_1_ column correspond to parameter combinations maximizing the *F*_1_ score as summarized in Table 5.

#### 3.2.3 False (FP & FN) predictions

Having assessed the best performing RippleNet model instance and combination of width and threshold parameters on the validation set (Table 5), we next analyze features FP and FN predictions on the validation dataset. LFP samples resulting in FP and/or FN predictions are illustrated in Figure 6-7. From this subset of samples, FP predictions appear to occur for transient events carrying power in the 150-250 Hz frequency range as reflected in both band-pass filtered LFPs (panels B) and LFP spectrograms (panels C), similar to correct (TP) predictions. One explanation may be that the procedure used to process the data initially either missed SPW-R events with poor signal-to-noise ratio, or that they were rejected manually based on some criteria. The prediction vectors *ŷ*^⟨*j*⟩^(*t*) approach a value of 1 in some FP cases, implying a high probability of an actual SPW-R event.

**Table 5:**
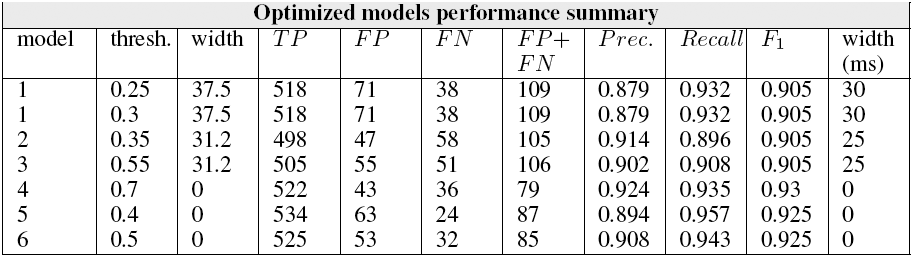
TP, FP, TN counts and performance metrics for different RippleNet instances on validation datasets, using optimized threshold settings.

**Figure 6:**
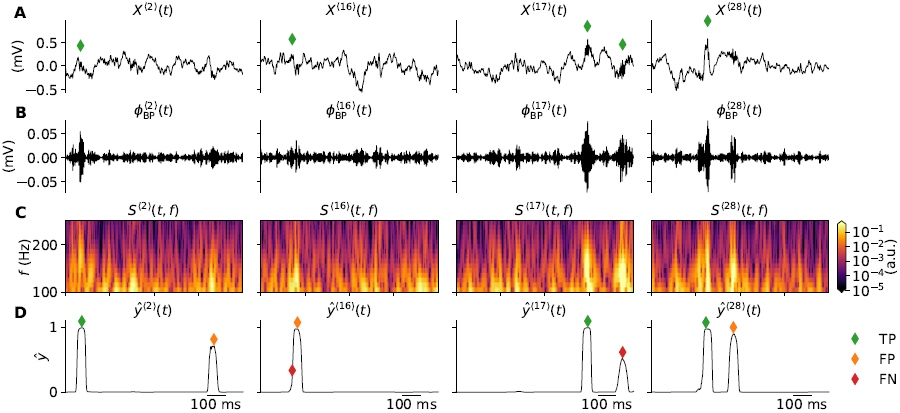
Examples of validation samples *j* resulting in FP and FN predictions. Each columns shows (A) input sequence, (B) bandpass-filtered LFP, (C) spectrograms and (D) predictions. The green, orange and red diamond markers denote times of TP, FP and FN events, respectively.

**Figure 7:**
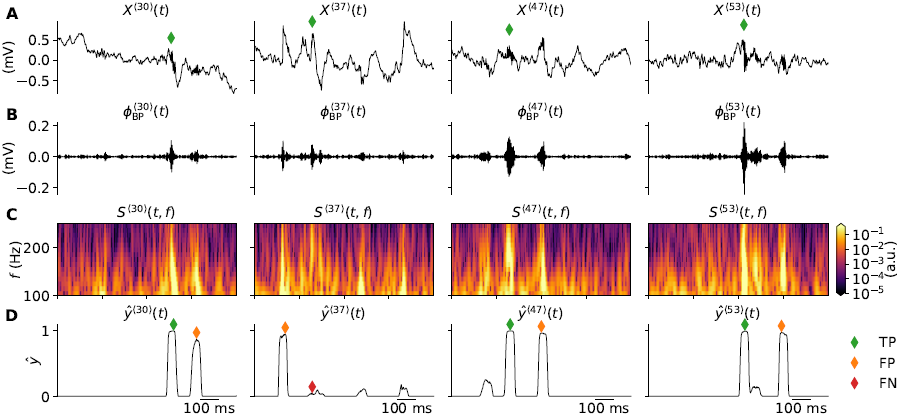
Same as Figure 6 showing another set of samples with detection mistakes.

For the set of samples resulting in FN predictions, the RippleNet make predictions *ŷ*(*t*) with magnitudes during the labeled SPW-R events that simply fail to produce a large enough amplitude and/or width for the peak-finding algorithm to detect the event. Here, a reduction of the threshold value for example will reduce FN counts, and increase FP and TP counts (cf. row 4 in Figure 5). Other cases resulting in both a FN and FP registration occurs if the predicted event time is outside of the boxcar shapes of the one-hot encoded signal. One such case is occurring in column 2 of Figure 6.

### 3.3 Ripple detection in time-continuous LFP data

With the same RippleNet instance as in the previous section, we next pay attention to the litmus test of this project, that is, applications to time continuous LFP recordings of arbitrary durations. We choose the 10 minute duration LFP signal of one session of one animal excluded from training or validation data (mouse 4029, session 1, see Table 1), and make predictions using RippleNet. This hold-out data set mimics new recordings unavailable at the time of training the networks. Predicted events within 1 s of movement periods are removed from the analysis to suppress FPs resulting from e.g., muscle noise.

By construction, the RippleNet algorithm can, in principle, be run on LFPs of arbitrary duration, even if all training and validation samples are of the same duration. We tested two operating modes: Either feeding in the entire LFP sequence as a single sample (not shown), or reshaping the LFP sequence into many sequential samples of the same duration. For the latter the predictions made on each sample (*ŷ*^⟨*j*⟩^(*t*)) can be concatenated together to form a continuous signal spanning the duration of the LFP entirely. In practice, 5-fold zero-padding and splitting of the signal into samples of duration 0.5 s, running predictions, concatenating predictions and computing the median output worked well on the hidden test set. Start-up transients in the output are suppressed by zero-padding the beginning and end of the full LFP signal by various amounts and realigning the predictions accordingly before computing the median.

For the 10 s segment *X*(*t*) shown in Figure 8A, with corresponding bandpass-filtered LFP, spectrogram and one-hot encoded events (panels B-D), all labeled SPW-R events are found (panel E). Not surprisingly, other significant responses with strengths above the peak-finding detection threshold are also found, resulting in a larger count of FPs compared to TPs (summarized in Table 6). Based on the previous analysis on a validation set with no negative samples that result in an error rate of about one per seven TP SPW-R event, we expect a higher frequency of FP predictions when predictions are made on samples spanning the entire session.

**Table 6:**
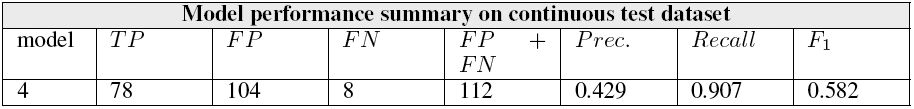
TP, FP, TN counts and performance metrics on continuous test set.

**Figure 8:**
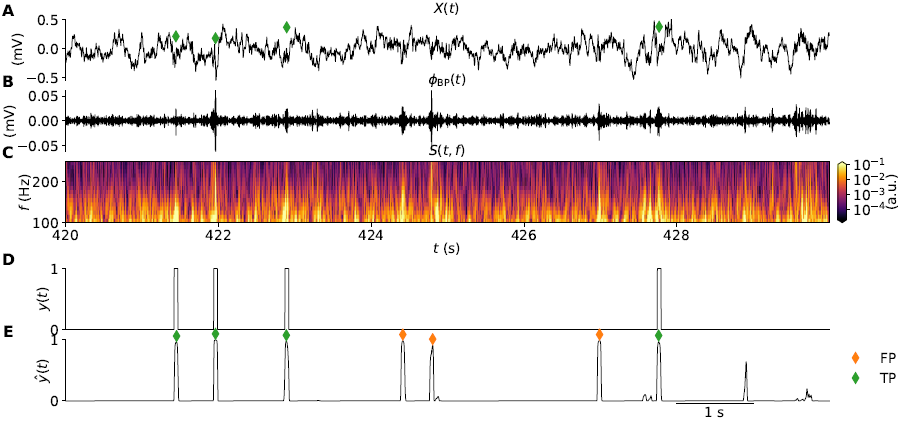
Application of RippleNet on continuous data. A: 10 s excerpt of input LFP signal *X*(*t*) = *ϕ*(*t*). The green diamonds marks the times of manually labeled SPW-R events. B: band-pass filtered LFP *ϕ*_BP_(*t*); C: Time-frequency resolved spectrogram *S*(*t, f*) of the LFP. D: label array *y*(*t*); E: prediction *ŷ*(*t*). The colored diamonds marks the times of predicted and missed SPW-R events using the threshold and width parameters that maximize the *F*_1_ score for the model.

For the approximate 10 minutes duration of the input LFP sequence, the chosen RippleNet instance finds about two times the number of events compared to the number of labeled SPW-R events in the input LFP sequence, see Figure 9A and Table 6. The cumulative count of predicted events appears linearly dependent on the cumulative count of labeled events. The cross-correlation coefficients *ρ*_*yŷ*_(*τ*) between predicted event times and labeled events in the test set (in bins of 2 ms) in Figure 9B, demonstrates a temporally precise prediction of event times, well within the 50 ms boxcar window around for each labeled SPW-R event in *y*(*t*) (Figure 8D).

**Figure 9:**
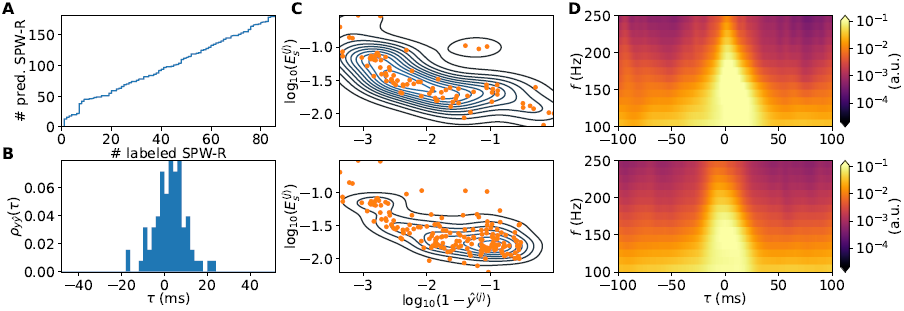
A: Cumulative counts of predicted SPW-R events as function of labeled SPW-R events. B: Cross-correlation coefficients between predicted ripple event times and labeled event times as function of time lag *τ* (2 ms bin size). C: Bandpass-filtered LFP SPW-R event energy 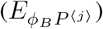 as function of (1 − *ŷ*^⟨*j*⟩^) of SPW-R events (orange dots). The contour lines show the bivariate kernel density estimate of the `kdeplot` method in the Seaborn plotting library. The top and bottom panel shows labeled and predicted SPW-R events, respectively. D: Averaged spectrograms for labeled (top) and predicted (bottom) SPW-R events.

#### 3.3.1 Features of predicted SPW-R events

Having established that the chosen RippleNet instance predicts presumed FP events at a high rate relative to TP rate in continuous LFP data, we next investigate the dependence between predicted SPW-R probability (*ŷ*^⟨*j*⟩^) and signal energy in the band-pass filtered LFP 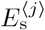 (Equation (14)). The RippleNet instance fares well with the labeled events in the hidden test set, with only a handful of FNs but many FPs (summarized in Table 6). The majority of labeled samples result in probabilities *ŷ*^⟨*j*⟩^ above the detection threshold 0.7. The ten samples with highest predicted probability are shown in Figure 10 rows 1-3, as well as the 10 samples with lowest predicted probability in rows 4-6. The RippleNet model instance recognizes SPW-R events with high amplitudes and quite stereotypical appearance both in the band-pass filtered LFP and spectrograms. At the lower end of the scale, SPW-R events show irregular fluctuations at lower amplitudes. The same holds true for the SPW-R events detected above threshold by the RippleNet algorithm (Figure 11). Detected events have transient activity around 150 Hz in their respective spectrograms.

**Figure 10:**
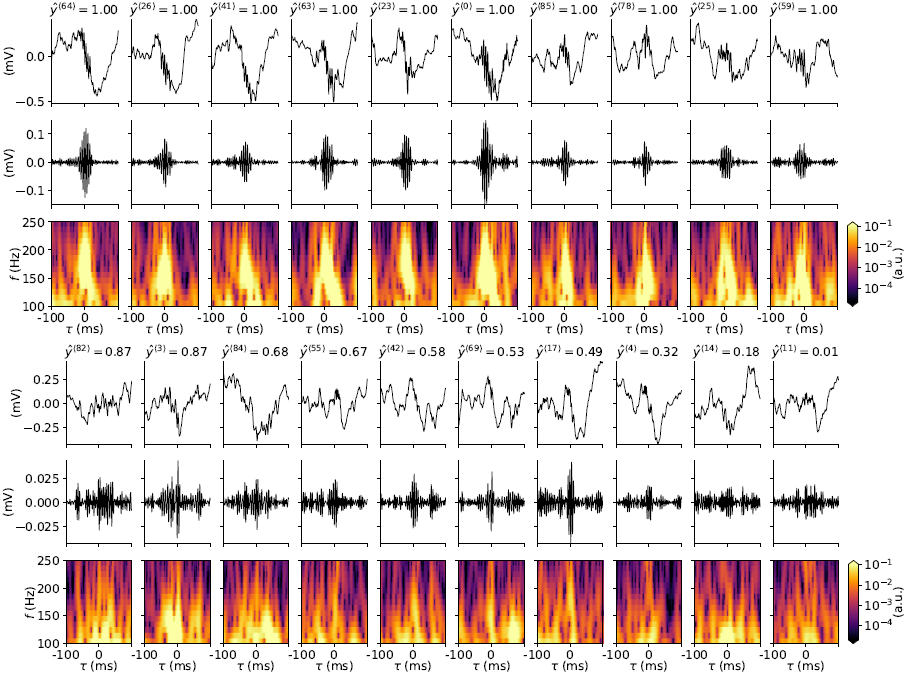
Labeled events (input LFP, bandpass-filtered signal, spectrograms) from the hidden test set with RippleNet confidence. Rows 1-3 shows events with high RippleNet-predicted probabilities (*ŷ*(*t*^⟨*j*⟩^) ≈ 1), rows 4-6 shows labeled 10 events with the lowest predicted event probability.

**Figure 11:**
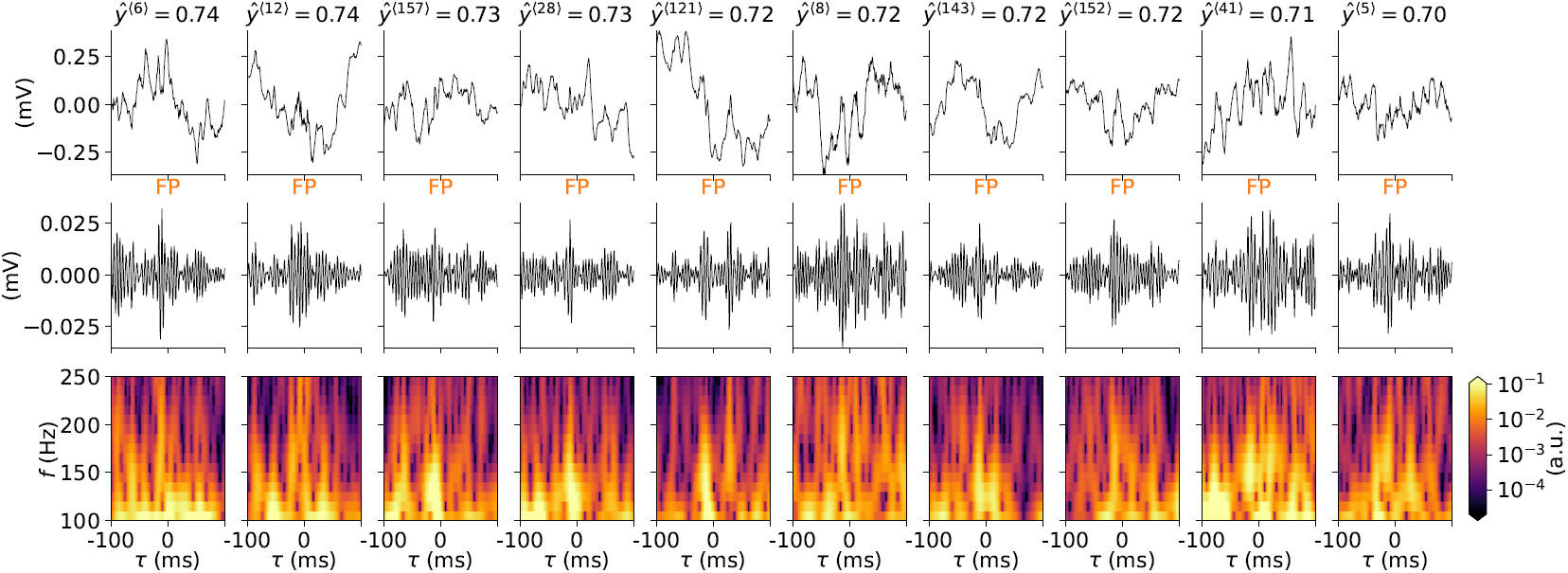
Same as Figure 10 but for 10 SPW-R events detected at or above threshold by the RippleNet algorithm. TP and FP status are shown in each column.

It thus appears that features of SPW-R events detected by the RippleNet algorithm share features of the manually scored events. In Figure 9C we plot the signal energy 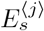 (Equation (14)) dependence on probability of *non*-event (1 − *ŷ*^⟨*j*⟩^) as predicted by the RippleNet instance. In this double-logarithmic plot, the distributions are overlapping, but many more RippleNet-detected events have lower energy and predicted probability. This finding is in agreement with the observed skewed distributions of SPW-R power (see e.g., Csicsvari et al. 1999a). We note that the averaged spectrogram of labeled and predicted events in Figure 9D are also very similar in appearance.

## 4 Discussion

### 4.1 Summary

In this paper we have introduced the RippleNet algorithm for detecting SPW-R events in time-continuous LFP data as recorded with single- or multi-channel probes in hippocampus CA1. The development of the RNN was motivated by high-performance speech recognition systems which utilize deep LSTM based RNNs (Graves, Mohamed, and Hinton 2013; Michalek and Vanek 2018). In the present context, the binary problem of detecting SPW-R events is simpler than speech recognition which must distinguish between different phonemes making up a spoken language. As such, the SPW-R detection task is analogous to mobile device wake up call detection to commands such as “Hello Siri!” or “OK, Google!” in noisy environments.

### 4.2 RippleNet performance on validation data

We trained different versions of RippleNet, each instantiated with different random weights on the same samples from the full set of manually scored data obtained in both mouse and rat CA1. On our validation dataset with mouse and rat data, the best-performing RippleNet instance resulted in 522 TP, 43 FP and 36 FN SWP-R predictions resulting in a combined *F*_1_ score of 0.93 (Table 5).

The non-causal versions of RippleNet utilizing bidirectional LSTMs is found to outperform the causal unidirectional versions during training and validation. On the validation data with optimized detection threshold settings, unidirectional RippleNets achieved similar TP counts, but with consistently higher error counts than bidirectional variants. The best-performing unidirectional RippleNet resulted in 518 TPs, 71 FPs and 38 FNs and *F*_1_ = 0.905.

The fact that the bidirectional versions outperform the unidirectional versions during training and validation, even if numbers of trainable parameters are larger for the latter cases is an indication that also the future context of the input LFP contains information about SPW-R events. Thus, real-time applications of RippleNet may be hampered by use of the unidirectional version, which may only use past and present context in order to make a prediction. For offline detection of SPW-R events the better choice is the bidirectional version.

### 4.3 RippleNet performance on test data

A hidden test dataset was also obtained from a single animal with one single session that is not included in any of the training/validation data. This best reflected real-world application to newly obtained LFP recordings in mouse. Features of actual SPW-R events may differ somewhat from those in the training/validation data obtained in different animals and species (Table 1). Test performance (in terms of loss *J, MSE, Precision, Recall, F*_1_) can be expected to be reduced compared to results obtained on the train and validation set. Indeed, the resulting counts of 78 TPs, 104 FPs and 8 FNs resulted in poor *Precision* (0.429) but still good *Recall* (0.907) and harmonic mean between the two (*F*_1_) of 0.582. We found that application of the RippleNet algorithm results in far more predictions of events with low energy than the conventional detection procedure used to label the test set initially.

When continuous LFP is used as input to the algorithm, RippleNet predicts many more events than were manually labeled in the test set, but superfluous events have similar features to labeled events. Given the nature of RNN parameters trained using backpropagation (Hochreiter and Schmidhuber 1997), the RippleNet algorithm may also be sensitive to hidden features in the LFP different than high-frequency (around 150 Hz or so) oscillations typically associated with SPW-R events, raising the question of whether or not conventional SPW-R detection algorithms relying on band-pass filtered LFPs discard useful information contained in other parts of the raw signal. One major caveat to the fact RippleNet algorithm finds most labeled events in the test and validation sets, but also many other positives, imply that the user must still make manual, quite likely subjective, judgements of whether or not detected events are true SPW-R events. As discussed below, results judged by a domain expert can be used to improve the method.

### 4.4 Improved data and extensions of RippleNet

#### Additional training data

Additional datasets containing labeled SPW-R events, available from online resources such as CRCNS.org (Teeters et al. 2008), can be added as soon as they become available. At present, several CRCNS deposits with CA1 LFPs have been made, but not every dataset comes with labeled SPW-R events. The uploaded datas are mostly obtained in rats using different kinds of electrodes such as laminar probes, tetrodes, and in different brain states such as sleep, anesthesized and awake states. While CA1 SPW-R events may represent underlying brain mechanisms that are highly preserved across species, it is *a priori* unclear if the SPW-R features our algorithm learned to recognize in the presently used mouse and rat datasets overlap with those in other data. The SPW-R events may for example have a different distributions of power across frequencies, or typical durations. For the present paper we opted to use only two sources of data, which should each be internally consistent in terms of data quality and methods (species, acquisition hardware, noise levels, data processing steps, label consistency etc.).

#### Data augmentation

Synthesizing recordings could also act as a potential supplement to real data. Generative Adverserial Nets (GAN) (Goodfellow et al. 2014) have for instance proven to produce very lifelike data in other domains such as image generation (e.g., Karras et al. 2019). There is an untapped potential to generate virtually unlimited amounts of ‘fake’ LFPs with similar statistics (power spectrum, temporal correlations, etc.) as the real data. A simple SPW-R model based on the superposition of modulated oscillatory events on pink (1*/f*) noise was already proposed by Sethi and Kemere 2014, but pure pink noise can not account for the temporal correlations of real data.

#### Neural network architecture

RNNs with LSTM layers or Gated Rectified Unit (GRU) layers (Chung et al. 2014) have for some time been considered state-of-the-art in sequence learning (Bai, Zico Kolter, and Koltun 2018). More recently, alternative architectures such as Temporal Convolutional Networks (TCN), for example WaveNet (van den Oord et al. 2016), also demonstrate capabilities of learning long-term temporal relationships in data. TCN networks were by Bai, Zico Kolter, and Koltun 2018 shown to outperform LSTM networks on various sequence learning tasks, and should also be evaluated for the SPW-R detection task described throughout this manuscript. The framework developed here around the high-level tensorflow.keras module allows for straight-forward comparison between different architectures. This comparison should also include conventional CNNs (LeCun, Bengio, and Hinton 2015) and variants such as deep residual networks (He et al. 2015) and inception networks (Szegedy et al. 2015; Ismail Fawaz et al. 2019). With the LSTM-based architectures we finally chose, one could potentially achieve even better performance by varying hyper parameters for the optimizer (e.g., learning rate), dropout layers (dropout rate), disabling batch-normalizing layers, optimize kernel sizes for the convolutional layers, add additional hidden layers and so forth.

#### Better-quality labels

While we here did not systematically compare predictions using fundamentally different architectures, we briefly tested multi-layered CNNs, causal TCNs, and replacing LSTM layers with GRU layers, but saw either worse or similar performance on the training and validation data. Similarly, we also tested increasing the layer sizes (and numbers of trainable parameters) and noted longer evaluation times and only slight improvements in accuracy. With the existing data and labels the features any deep learning method may learn is limited if labels are inaccurate. For instance, the rat dataset contained information on SPW-R durations (which we ignored) while the mouse data only contained their locations. More accurate predictions on the existing dataset during training and validation can be achieved by more thorough labeling, perhaps by multiple experts independently.

#### Self-improving method

As soon as RippleNet has been used to find SPW-R event times in batches of new data, validated SPW-R events can supplement the initial training dataset. Then, the pre-trained RippleNet instance presented here can be trained for more iterations, learn new features and consolidate learned features present in the initial and new samples. With time and several such iterations, an even better performance can be achieved. Another possibility is classification of different kinds of SPW-R events and non-SPW-R (noise) events. We did not yet persuade such classification, as it would require a modification to the final dense output layer to use the so-called Softmax activation function instead of the presently used sigmoid activation function. The output dimensionality and values would then reflect the number of classes and respective probabilities.

#### Online and offline RippleNet applications

The bidirectional LSTM architecture is here found to perform better than the causal, unidirectional LSTM variant. Only the latter may see potential use in online and real-time applications, which only rely on past and present segments of the input time series. High temporal accuracy may be achieved, and unidirectional RippleNet instances may therefore see potential use in closed loop experiments where a stimulus may be delivered at the times of detected SPW-R events. As the RippleNet algorithm runs on temporally downsampled LFP signals, a good realtime factor is achievable on short segments, in particular if the computer has GPU accelerating capabilities.

Pre-trained RippleNet models can easily be loaded (with the tf.keras.load_model function in Python), and can be incorporated into Python-based data processing workflows with ease. For this purpose, models may also be converted to the higher-level tf.estimator API.

#### Web/cloud applications

The majority of development and analysis of RippleNet was incorporated using Jupyter notebooks^14^ running on the Google Colaboratory portal^15^ with data file access and synchronization via Google Drive^16^. RippleNet can be provided as a Cloud service, or as a service running locally on the user’s computer. The latter may facilitate on not having to upload potentially large files but will greatly benefit from a local GPU in order to accelerate compute times. Distribution of RippleNet to end users can be done using containers in Docker^17^, Kubernetes^18^ or similar. A cloud service however would facilitate on the powerful GPU backends provided via services like Google Cloud^19^ which also has efficient data handling.

Another option could also be a port of RippleNet to tensorflow.js as the model is already only using keras constructs. The conversion step appears trivial^20^ and could allow execution of RippleNet in html contexts.

#### Graphical User Interface (GUI)

The current RippleNet version is set up as a step-by-step workflows in Jupyter notebooks for training, validation and application to continuous data, respectively. While a standalone, interactive RippleNet application with a GUI is certainly possible to develop using cross-platform application tools such as PyQT^21^, it is presently only considered. A prototype jupyter notebook which allows for user-interactive rejection of detected events (noise events) and storage of accepted events has been implemented, however.

### 4.5 Outlook

This work is an effort to introduce novel machine learning and deep learning algorithms for the detection of SPW-R events in electrophysiological data. The RippleNet algorithm presented here learns through supervised learning an internal representation of features of SPW-R events, which facilitates a highly non-linear transformation of input LFP signals into output signals that represent the time-varying probabilities of SPW-R events. This represents a fundamental change from the typical procedure employed in standard detection workflows relying on hand crafted feature extraction. With access to more training data with labeled events, the method can improve by running more training iterations on new data. Our hope is that the tool can reduce the amount of time the experimentalist spend on manual extraction of SPW-R events using heuristic criteria, and allow for a better understanding of features of these events and their role in brain function. Also, we hope that this powerful framework may be adapted to other detection tasks, for instance onset of epileptic seizures. In this direction, there are many potential applications, also clinical.

## 5 Data and code availability statement

All source codes and data to reproduce the findings and illustrations of this paper are available at github.com/espenhgn/RippleNet and Zenodo.org (Hagen 2020).

## 6 Author contributions

EH, KHP, RE and AJS conceived and conceptualized the project. EH wrote the paper. EH, ARC, RE, KHP, GTE and AJS cowrote the paper. ARC and RE measured mouse LFP datas and labeled SPW-R events. AJS wrote the manual SPW-R detection code. EH wrote and executed all codes for extracting training/validation/test datasets, neural networks, neural-network training, analysis and plots for this paper.

## 7 Funding

This work was funded by the Research Council of Norway (NFR) through the grant/award numbers 250128 (COBRA), 300504 (IKTPLUSS), 274328, 249988 and NS9021K (NIRD), the Marie Skłodowska-Curie IF 753608, and the Letten Foundation.

## 8 Acknowledgements

We would like to thank the Buzsaki lab (buzsakilab.com) and David Tingley in particular for publicly sharing their valuable electrophysiological datasets at https://buzsakilab.com/wp/datasets/.

Finally we would like to thank Google for facilitating free access to compute resources and storage through the Google Colaboratory portal and Google Drive.

## Footnotes

1 https://www.mathworks.com

2 https://buzsakilab.com/wp/datasets

3 python.org

4 jupyter.org

5 numpy.org

6 scipy.org

7 h5py.org

8 matplotlib.org

9 pandas.pydata.org

10 www.anaconda.com

11 seaborn.pydata.org

12 tensorflow.org

13 colab.research.google.com

14 jupyter.org

15 colab.research.google.com

16 drive.google.com

17 docker.com

18 kubernetes.io

19 cloud.google.com

20 tensorflow.org/js/tutorials/conversion/import_keras

21 riverbankcomputing.com/static/Docs/PyQt5

## Notes

https://github.com/espenhgn/RippleNet

### Competing Interest Statement

The authors have declared no competing interest.

## References

Abadi, Martín et al. (2015). TensorFlow: Large-Scale Machine Learning on Heterogeneous Systems. Software available from tensorflow.org. URL: https://www.tensorflow.org/.

Axmacher, Nikolai, Christian E Elger, and Juergen Fell (May 2008). “Ripples in the medial temporal lobe are relevant for human memory consolidation.” In: Brain 131.7 (Pt 7), pp. 1806–1817. ISSN: 1460-2156. DOI: 10.1093/brain/awn103.

Bai, Shaojie, J. Zico Kolter, and Vladlen Koltun (Mar. 2018). “An Empirical Evaluation of Generic Convolutional and Recurrent Networks for Sequence Modeling”. In: arXiv e-prints, 1803.01271. arXiv: 1803.01271 [cs.LG].

Buzsaki, G et al. (May 1992). “High-frequency network oscillation in the hippocampus”. In: Science 256.5059 (5059), pp. 1025–1027. ISSN: 0036-8075. DOI: 10.1126/science.1589772.

Buzsáki, G. et al. (Jan. 2003). “Hippocampal network patterns of activity in the mouse”. In: Neuroscience 116.1 (1), pp. 201–211. ISSN: 0306-4522. DOI: 10.1016/s0306-4522(02)00669-3.

Buzsáki, György (Sept. 2015). “Hippocampal sharp wave-ripple: A cognitive biomarker for episodic memory and planning”. In: Hippocampus 25.10 (10), pp. 1073–1188. ISSN: 1098-1063. DOI: 10.1002/hipo.22488.

Buzsáki, György, Nikos Logothetis, and Wolf Singer (Oct. 2013). “Scaling Brain Size, Keeping Timing: Evolutionary Preservation of Brain Rhythms”. In: Neuron 80.3 (3), pp. 751–764. ISSN: 1097-4199. DOI: 10.1016/j.neuron.2013.10.002.

Caputi, Antonio et al. (2012). “Selective Reduction of AMPA Currents onto Hippocampal Interneurons Impairs Network Oscillatory Activity”. In: PLoS ONE 7.6 (6). Ed. by Huibert D. Mansvelder, e37318. ISSN: 1932-6203. DOI: 10.1371/journal.pone.0037318.

Chung, Junyoung et al. (Dec. 2014). “Empirical Evaluation of Gated Recurrent Neural Networks on Sequence Modeling”. In: arXiv e-prints, 1412.3555. arXiv: 1412.3555 [cs.NE].

Collette, Andrew et al. (2019). h5py/h5py: 2.10.0. DOI: 10.5281/ZENODO.3401726.

Csicsvari, Jozsef et al. (Aug. 1999a). “Fast Network Oscillations in the Hippocampal CA1 Region of the Behaving Rat”. In: The Journal of Neuroscience 19.16 (16), RC20–RC20. ISSN: 1529-2401. DOI: 10.1523/jneurosci.19-16-j0001.1999.

Csicsvari, Jozsef et al. (Jan. 1999b). “Oscillatory Coupling of Hippocampal Pyramidal Cells and Interneurons in the Behaving Rat”. In: The Journal of Neuroscience 19.1 (1), pp. 274–287. ISSN: 0270-6474. DOI: 10.1523/jneurosci.19-01-00274.1999.

Csicsvari, Jozsef et al. (Nov. 2000). “Ensemble Patterns of Hippocampal CA3-CA1 Neurons during Sharp Wave–Associated Population Events”. In: Neuron 28.2 (2), pp. 585–594. ISSN: 0896-6273. DOI: 10.1016/s0896-6273(00)00135-5.

Davidson, Thomas J., Fabian Kloosterman, and Matthew A. Wilson (Aug. 2009). “Hippocampal Replay of Extended Experience”. In: Neuron 63.4 (4), pp. 497–507. ISSN: 1097-4199. DOI: 10.1016/j.neuron.2009.07.027.

Eggermont, Jos J. (2010). “Analysis of Parallel Spike Trains”. In: ed. by Stefan Rotter Sonja Grün. Vol. 7. Springer Series in Computational Neuroscience. Springer US. Chap. 5, pp. 77–102. ISBN: 978-1-4419-5674-3. DOI: 10.1007/978-1-4419-5675-0.

Einevoll, Gaute T. et al. (2013). “Modelling and analysis of local field potentials for studying the function of cortical circuits”. In: Nature Reviews Neuroscience 14.11 (11), pp. 770–785. ISSN: 1471-0048. DOI: 10.1038/nrn3599.

Fawaz, Hassan Ismail et al. (2019). “Deep learning for time series classification: a review”. In: Data Mining and Knowledge Discovery 33.4, pp. 917–963. DOI: 10.1007/s10618-019-00619-1.

Fritsch, C., A. Ibanez, and M. Parrilla (Dec. 1999). “A digital envelope detection filter for real-time operation”. In: IEEE Transactions on Instrumentation and Measurement 48.6, pp. 1287–1293. ISSN: 1557-9662. DOI: 10.1109/19.816150.

Goodfellow, Ian J. et al. (June 2014). “Generative Adversarial Networks”. In: arXiv e-prints, 1406.2661. arXiv: 1406.2661 [stat.ML].

Graves, Alex, Abdel-rahman Mohamed, and Geoffrey Hinton (Mar. 2013). “Speech Recognition with Deep Recurrent Neural Networks”. In: arXiv e-prints, 1303.5778. arXiv: 1303.5778 [cs.NE].

Hagen, Espen (May 2020). espenhgn/RippleNet: RippleNet-v0.1. Version v0.1. DOI: 10.5281/ZENODO.3819821. URL: https://doi.org/10.5281/zenodo.3819821.

Hagen, Espen et al. (Oct. 2016). “Hybrid Scheme for Modeling Local Field Potentials from Point-Neuron Networks”. In: Cerebral Cortex 26.12, pp. 4461–4496. ISSN: 1460-2199. DOI: 10.1093/cercor/bhw237. URL: https://doi.org/10.1093/cercor/bhw237.

He, Kaiming et al. (Dec. 2015). “Deep Residual Learning for Image Recognition”. In: arXiv e-prints, 1512.03385. arXiv: 1512.03385 [cs.CV].

Hochreiter, Sepp and Jürgen Schmidhuber (Nov. 1997). “Long Short-Term Memory”. In: Neural Computation 9.8 (8), pp. 1735–1780. ISSN: 0899-7667. DOI: 10.1162/neco.1997.9.8.1735.

Hunter, J. D. (2007). “Matplotlib: A 2D Graphics Environment”. In: Computing in Science Engineering 9.3, pp. 90–95.

Ismail Fawaz, Hassan et al. (Sept. 2019). “InceptionTime: Finding AlexNet for Time Series Classification”. In: arXiv e-prints, 1909.04939. arXiv: 1909.04939 [cs.LG].

Jadhav, S. P. et al. (May 2012). “Awake Hippocampal Sharp-Wave Ripples Support Spatial Memory”. In: Science 336.6087 (6087), pp. 1454–1458. ISSN: 1095-9203. DOI: 10.1126/science.1217230.

James Conder (2020). gaussfilt(t,z,sigma). https://www.mathworks.com/matlabcentral/fileexchange/43182-gaussfilt-t-z-sigma. Retrieved March 30, 2020.

John O’Keefe, Lynn Nadel (1978). The hippocampus as a cognitive map. Oxford University Press, Oxford. ISBN: 0-19-857206-9.

Karras, Tero et al. (Dec. 2019). “Analyzing and Improving the Image Quality of StyleGAN”. In: arXiv e-prints, 1912.04958. arXiv: 1912.04958 [cs.CV].

Kingma, Diederik P. and Jimmy Ba (Dec. 2014). “Adam: A Method for Stochastic Optimization”. In: arXiv e-prints, 1412.6980. arXiv: 1412.6980 [cs.LG].

LeCun, Yann, Yoshua Bengio, and Geoffrey Hinton (May 2015). “Deep learning”. In: Nature 521.7553 (7553), pp. 436–444. ISSN: 1476-4687. DOI: 10.1038/nature14539.

MATLAB (2018). version 9.5.0.1067069 (R2018b) Update 4. Natick, Massachusetts.

McKinney, Wes (2010). “Data Structures for Statistical Computing in Python”. In: Proceedings of the 9th Python in Science Conference. Ed. by Stéfan van der Walt and Jarrod Millman. SciPy, pp. 56 –61. DOI: 10.25080/majora-92bf1922-00a.

Michalek, Josef and Jan Vanek (June 2018). “A Survey of Recent DNN Architectures on the TIMIT Phone Recognition Task”. In: arXiv e-prints, arXiv:1806.07974. arXiv: 1806.07974 [cs.CL].

Norman, Yitzhak et al. (Aug. 2019). “Hippocampal sharp-wave ripples linked to visual episodic recollection in humans.” In: Science 365.6454 (6454), eaax1030. ISSN: 1095-9203. DOI: 10.1126/science.aax1030.

Petersen, Peter Christian, Michelle Hernandez, and György Buzsáki (2018). Public electrophysiological datasets collected in the Buzsaki Lab. en. DOI: 10.5281/ZENODO.3629881.

Ramirez-Villegas, Juan F., Nikos K. Logothetis, and Michel Besserve (Nov. 2015). “Diversity of sharp-wave–ripple LFP signatures reveals differentiated brain-wide dynamical events”. In: Proceedings of the National Academy of Sciences 112.46 (46), E6379–E6387. ISSN: 1091-6490. DOI: 10.1073/pnas.1518257112.

Rawat, Waseem and Zenghui Wang (Sept. 2017). “Deep Convolutional Neural Networks for Image Classification: A Comprehensive Review.” In: Neural Computation 29.9 (9), pp. 2352–2449. ISSN: 1530-888X. DOI: 10.1162/neco_a_00990.

Schomburg, E. W. et al. (Aug. 2012). “The Spiking Component of Oscillatory Extracellular Potentials in the Rat Hippocampus”. In: The Journal of Neuroscience 32.34 (34), pp. 11798–11811. ISSN: 1529-2401. DOI: 10.1523/jneurosci.0656-12.2012.

Sethi, Ankit and Caleb Kemere (Aug. 2014). “Real time algorithms for sharp wave ripple detection”. In: Conf Proc IEEE Eng Med Biol Soc 2014, pp. 2637–2640. ISSN: 1557-170X. DOI: 10.1109/embc.2014.6944164.

Silva, Fernando Lopes da (Dec. 2013). “EEG and MEG: Relevance to Neuroscience”. In: Neuron 80.5 (5), pp. 1112–1128. ISSN: 1097-4199. DOI: 10.1016/j.neuron.2013.10.017.

Szegedy, Christian et al. (June 2015). “Going deeper with convolutions”. In: 2015 IEEE Conference on Computer Vision and Pattern Recognition (CVPR). IEEE, pp. 1–9. DOI: 10.1109/cvpr.2015.7298594.

Teeters, Jeffrey L. et al. (Feb. 2008). “Data Sharing for Computational Neuroscience”. In: Neuroinformatics 6.1 (1), pp. 47–55. ISSN: 1559-0089. DOI: 10.1007/s12021-008-9009-y.

Tingley, David and György Buzsáki (June 2018). “Transformation of a Spatial Map across the Hippocampal-Lateral Septal Circuit”. In: Neuron 98.6 (6), 1229–1242.e5. ISSN: 1097-4199. DOI: 10.1016/j.neuron.2018.04.028.

Tingley, David (Jan. 2020). “Routing of Hippocampal Ripples to Subcortical Structures via the Lateral Septum”. In: Neuron 105.1 (1), 138–149.e5. ISSN: 1097-4199. DOI: 10.1016/j.neuron.2019.10.012.

van den Oord, Aaron et al. (Sept. 2016). “WaveNet: A Generative Model for Raw Audio”. In: arXiv e-prints, 1609.03499. arXiv: 1609.03499 [cs.SD].

van der Walt, S., S. C. Colbert, and G. Varoquaux (2011). “The NumPy Array: A Structure for Efficient Numerical Computation”. In: Computing in Science Engineering 13.2, pp. 22–30.

Virtanen, Pauli et al. (2020). “SciPy 1.0: Fundamental Algorithms for Scientific Computing in Python”. In: Nature Methods 17, pp. 261–272. DOI: https://doi.org/10.1038/s41592-019-0686-2.

Waskom, Michael et al. (2020). mwaskom/seaborn: v0.10.1 (April 2020). DOI: 10.5281/ZENODO.3767070.

Zuo, Rui et al. (Feb. 2019). “Automated Detection of High-Frequency Oscillations in Epilepsy Based on a Convolutional Neural Network”. In: Frontiers in Computational Neuroscience 13, p. 6. ISSN: 1662-5188. DOI: 10.3389/fncom.2019.00006.

